# Alzheimer’s patient brain myeloid cells exhibit enhanced aging and unique transcriptional activation

**DOI:** 10.1101/610345

**Authors:** Karpagam Srinivasan, Brad A. Friedman, Ainhoa Etxeberria, Melanie A. Huntley, Marcel P. van der Brug, Oded Foreman, Jonathan S. Paw, Zora Modrusan, Thomas G. Beach, Geidy E. Serrano, David V. Hansen

## Abstract

Gene expression changes in brain microglia from mouse models of Alzheimer’s disease (AD) are highly characterized and reflect specific myeloid cell activation states that could modulate AD risk or progression. While some groups have produced valuable expression profiles for human brain cells^1–4^, the cellular clarity with which we now view transcriptional responses in mouse AD models has not yet been realized for human AD tissues due to limited availability of fresh tissue samples and technological hurdles of recovering transcriptomic data with cell-type resolution from frozen samples. We developed a novel method for isolating multiple cell types from frozen post-mortem specimens of superior frontal gyrus for RNA-Seq and identified 66 genes differentially expressed between AD and control subjects in the myeloid cell compartment. Myeloid cells sorted from fusiform gyrus of the same subjects showed similar changes, and whole tissue RNA analyses further corroborated our findings. The changes we observed did not resemble the “damage-associated microglia” (DAM) profile described in mouse AD models^5^, or other known activation states from other disease models. Instead, roughly half of the changes were consistent with an “enhanced human aging” phenotype, whereas the other half, including the AD risk gene *APOE*, were altered in AD myeloid cells but not differentially expressed with age. We refer to this novel profile in <u>h</u>uman <u>A</u>lzheimer’s <u>m</u>icroglia/myeloid cells as the HAM signature. These results, which can be browsed at research-pub.gene.com/BrainMyeloidLandscape/reviewVersion, highlight considerable differences between myeloid activation in mouse models and human disease, and provide a genome-wide picture of brain myeloid activation in human AD.

## Main Text

We recently reported a unique method for isolating multiple cell types from fresh mouse brain, involving mechanical dissociation at 4ºC, ethanol fixation, immunolabeling, and FACS followed by RNA-Seq^6^. We next tested whether this approach (and others; see Methods) was suitable for collecting various cell populations from frozen human brain tissues. When analysis of cell type markers by qPCR confirmed the desired levels of cell type enrichment and purity, we then used the method to collect neuronal (NeuN^+^), astrocytic (GFAP^+^), endothelial (CD31^+^), and myeloid (CD11b^+^) cell bodies (DAPI^+^) from superior frontal gyrus (SFG) of 22 AD (Braak stage V-VI) and 21 neurologically normal age-, sex- and post-mortem interval-matched controls, followed by RNA purification and sequencing (Fig. 1a, Extended Data Fig. 1, Supplementary Table 1). Due to methodological limitations (e.g., use of frozen tissues; see Methods), we had to discard a number of unacceptable RNA profiles based on principal component analysis (PCA), cell type marker, and other QC analyses (Extended Data Fig. 2), yielding a total of 115 cell type-specific expression profiles, including myeloid profiles from 15 control and 10 AD subjects (Fig. 1b, c).

**Figure 1.**
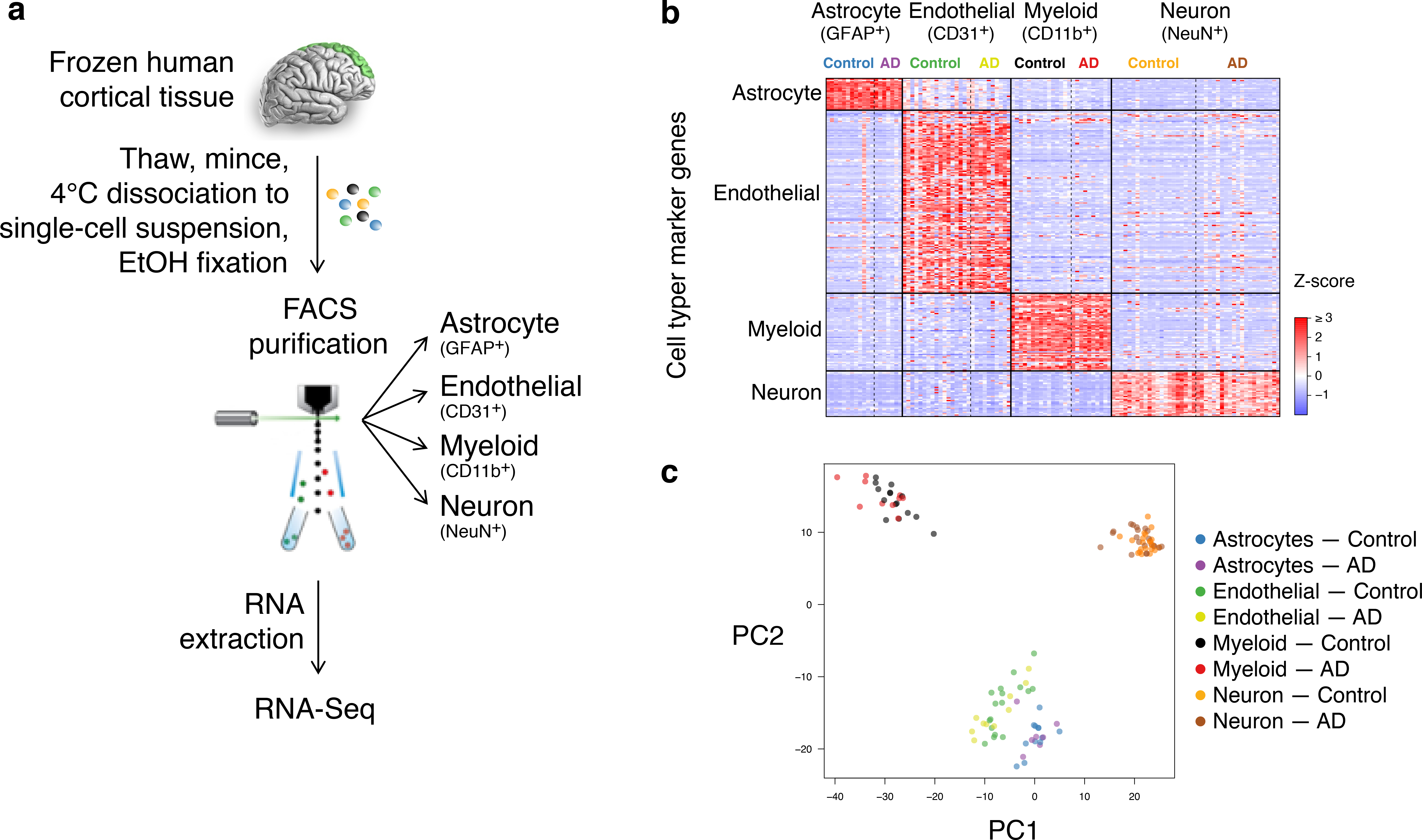
Expression profiling of sorted cell populations from frozen post-mortem human brain tissue. **(a)** Experimental overview. **(b)** Expression of known cell-type markers, derived from previously published fresh-sorted human cells ^4^, in QC-passing expression profiles, indicates high cell type purity. Each gene was Z-score normalized across all profiles of all cell types and values were plotted in a heatmap. **(c)** Principal components analysis using most variable genes reveals good separation of four cell types.

**Figure 2.**
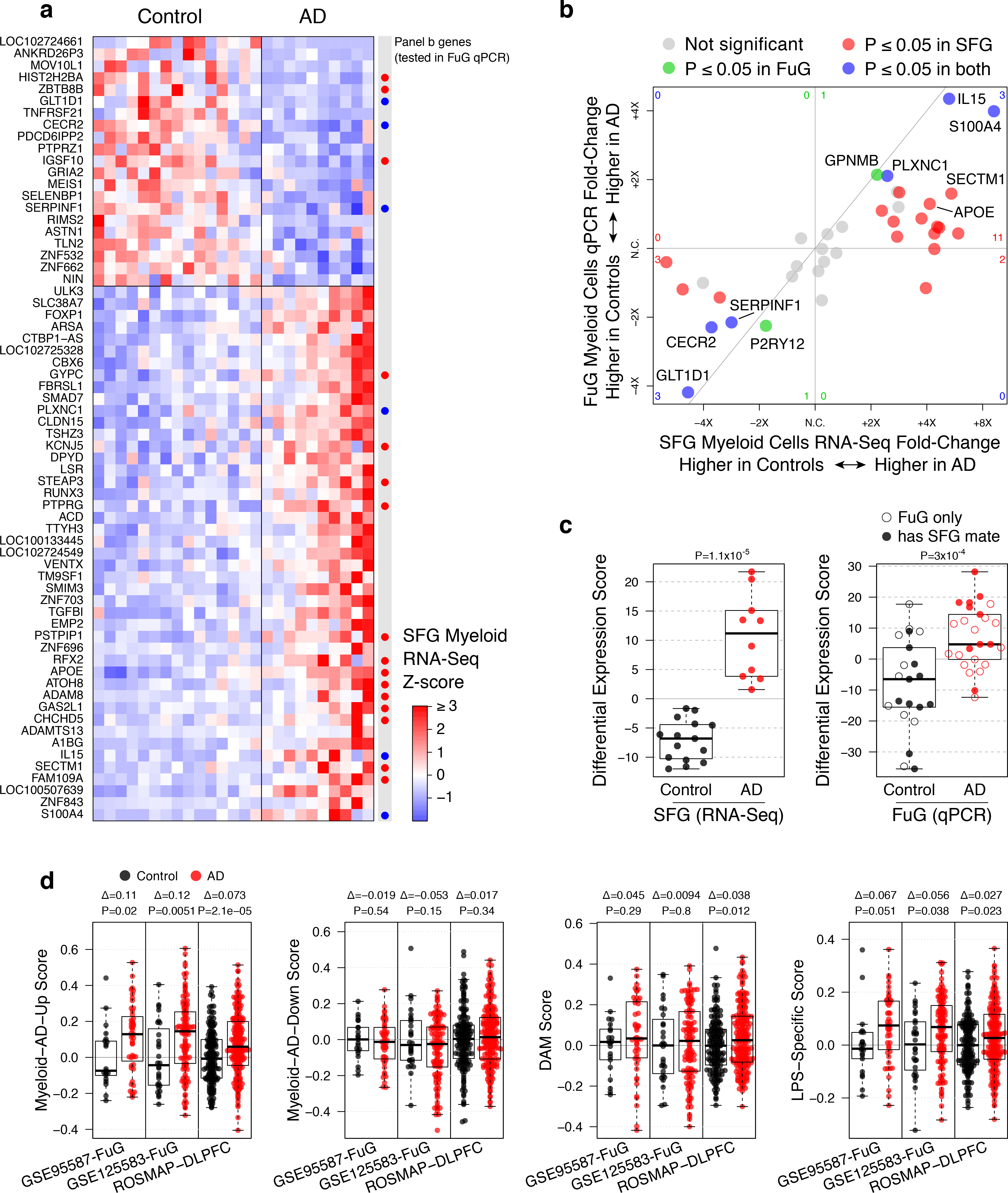
Human myeloid cells exhibit an AD-associated differential expression (DE) profile. **(a)** Heatmap of AD-differentially expressed genes (rows, DESeq2 adjusted *P* ≤ 0.05 and maximum Cook’s P ≥ 0.01) in control and AD SFG-derived myeloid expression profiles (columns). Columns are sorted by diagnosis then first principal component of Z-score matrix. “Panel B Genes” indicates genes that were subsequently assayed by qPCR in FuG sorted myeloid cells, with colors from panel B. **(b)** “4-way” comparing AD-associated DE in SFG measured by RNA-Seq (x-axis) to DE in FuG measured by qPCR (*y*-axis). Each point represents one gene colored by whether adjusted *P*-value was ≤ 0.05 in one or both of the DE analysis. Genes were selected manually, consisting of about 1/3 of the DE genes from the RNA-Seq study and several other cell type markers and genes of interest. **(c)** SFG DE is reproduced in FuG. DE scores (methods) are shown for each SFG and FuG sample, using the SFG DE genes that were included in the qPCR panel. For FuG samples, open circles indicate that a QC-passing SFG RNA-Seq profile was not available from that subject. *P* value, *t*-test. **(d)** Human myeloid AD expression changes are recapitulated in myeloid-balanced bulk AD tissues, and are more robustly altered than mouse-derived myeloid changes. Each study was separately myeloid-balanced to create a subset of samples with similar myeloid gene set scores, and neuronal genes were removed from each gene set (all samples and genes shown in Extended Data Fig. 4c). Each panel shows gene set scores for the indicated gene sets for each of the myeloid-balanced AD or control samples. Δ, mean log2-fold change; *P*-value, t-test

Differential expression (DE) analysis using DESeq2 identified 45 genes increased and 21 decreased in AD myeloid cells relative to controls (at adjusted *P*-value ≤ 0.05, and maximum Cook’s *P*-value ≥ 0.01 to help exclude hits driven by outliers; Fig. 2a; see Methods). To assess the robustness of these findings, we used the same method to isolate cell types from frozen fusiform gyrus (FuG) of the same subjects plus others, totaling 25 AD and 21 control tissues, and quantified gene expression in the myeloid cells using qPCR for genes of interest including a subset of SFG DE genes (16 up genes, 6 down genes). The direction of effect across samples was replicated for nearly every DE gene tested (Fig. 2b), and when we used these genes to assign a DE score to each tissue sample we again saw a clear difference between AD myeloid cells and controls (Fig. 2c). Moreover, for subjects with both SFG and FuG data available, the DE scores were correlated between the two regions (Extended Data Fig. 3a).

**Figure 3.**
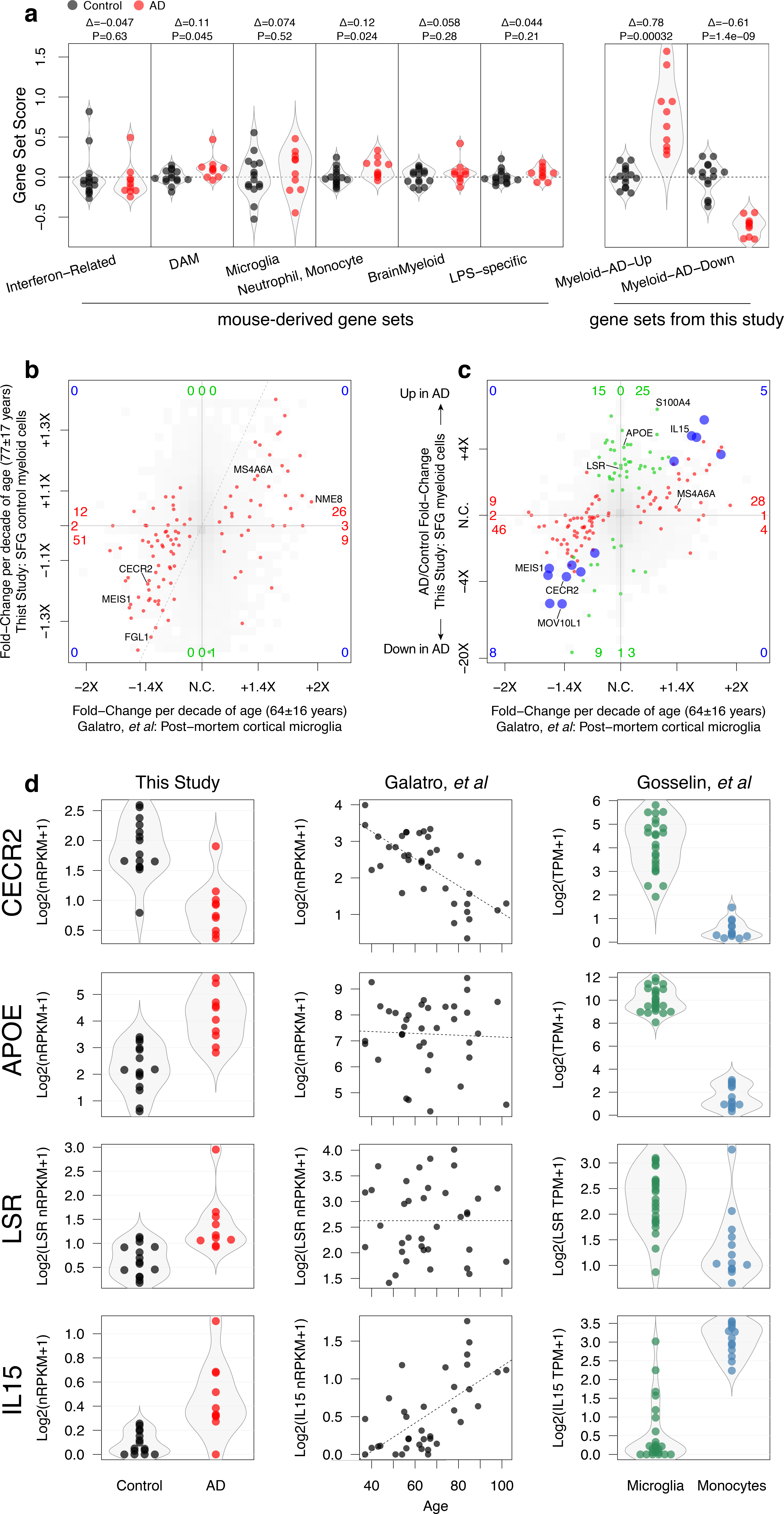
Human AD Myeloid (HAM) gene expression changes share similarities with normal aging. **(a)** Distribution of scores for mouse- and human-derived gene sets in SFG myeloid cell profiles. *P* values, *t*-test. Δ, mean log2-fold change. **(b)** Previously reported normal age-associated DE is recapitulated in control subjects of this study. 4-way plot (like Fig. 2b) shows age-associated DE from Galatro *et al* on *x*-axis and from this study’s controls on *y*-axis. Genes in red met adjusted *P* ≤ 0.05 cutoff in Galatro; other genes shown as smoothed density in shades of gray. No DE genes from Galatro met *P* ≤ 0.05 cutoff for aging in our study, but most trended in a consistent direction (bottom left and top right quadrants). The lack of statistical significance and muted fold-changes in our study may result from having fewer samples and a more limited age range. *(c)* AD myeloid cells exhibit accelerated aging and a new type of transcriptional activation. 4-way plot shows same *x*-axis as panel b; *y*-axis shows DE between AD and control myeloid cells. Color indicates *P* ≤ 0.05 significance with aging only (red), with AD only (green) or with both (blue). Most “red” genes, DE with age, trended in a consistent direction with AD (bottom left and top right quadrants), indicating that AD myeloid cells exhibit “enhanced aging”. The “green” genes near the top, including *APOE*, indicate an AD-specific signature that is distinct from normal aging. **(d)** Example gene expression plots. Each point shows the expression of the indicated gene in a single sample in one of the three studies. In middle column (Galatro *et al*) dashed line indicates best linear fit.

Another way to validate our DE results was to examine whole tissue RNA datasets from AD and control patients. Despite their limitations^6^, such datasets allow the evaluation of larger cohorts. We examined three studies: our previously published cohort of FuG samples (GSE95587), a newly generated FuG cohort (GSE125583) (Extended Data Fig. 4a), and the ROSMAP cohort^7^ from the dorsolateral prefrontal cortex (DLPFC). We used “myeloid-balancing” (involving reductions in sample size) to control for differences in myeloid content between control and AD tissues (Extended Data Fig. 4b), excluded neuronal-enriched genes from the gene sets to mitigate the confounding effects of neuronal loss, and then calculated gene set scores for the AD-Up and AD-Down genes we observed in SFG myeloid cells (Fig. 2d). In all three whole tissue datasets, the Myeloid-AD-Up gene set was significantly increased. This increase was more apparent in later Braak stages, which also showed decreased expression of our Myeloid-AD-Down genes set (Extended Data Fig. 4c). (The Braak stage analyses did not include the corrective measures for altered cellular makeup of AD tissues since it was impractical to reduce the sample size.) These whole tissue RNA analyses provided additional evidence that our DE findings in AD myeloid cells sorted from SFG and FuG were present in multiple regions of cortex.

We recently used microglial expression data from diverse mouse models to define gene modules related to various activation states, and we reported that expression signals for some of these modules were slightly elevated in whole tissue RNA profiles from AD brains^8^. Compared to these mouse-derived gene sets, our Myeloid-AD-Up gene set was more robustly elevated in AD whole tissue RNA (Fig. 2d, Extended Data Fig. 4c). We wondered whether expression of the mouse-derived gene sets might be more apparent in our isolated human myeloid cells than in whole tissue, but any such AD-related changes were again subtle if at all present, especially compared to the AD-Up and AD-Down gene sets defined herein (Fig. 3a). In fact, out of more than a hundred neurodegeneration-related/DAM genes identified in mouse, only one was significantly increased in SFG microglia from AD patients—APOE (4.1-fold change, P = 0.00040) (Extended Data Fig. 5a). Interestingly, the DAM gene with the next closest *P*-value was *APOC1* (3.0-fold change, P = 0.056). *GPNMB* also showed a strong upward trend (2.2-fold change, P = 0.11), but most DAM genes’ upward (n=58) or downward (n=54) trends were weak. For example, *ITGAX*, *CCL3*, and *CXCR4* had fold changes of 1.2, 0.84, and 0.80, with *P*-values of 0.82, 0.90, and 0.88, respectively.

Perhaps equally surprising, the “Brain Myeloid” gene sets that define resting or “homeostatic” microglia and are downregulated in response to virtually any perturbation in mice^8^ showed no hint of downregulation in our SFG myeloid cells sorted from AD brains (Fig. 3a). Of over a hundred genes in this set, only *SERPINF1* (fold change = 0.35, P = 0.0062) showed the significant reduction predicted by mouse data (Extended Data Fig. 5b, c). *P2RY12* (fold change = 0.54, P = 0.50) and 77 other genes trended downward, while *CX3CR1* (fold change = 1.04, P = 0.96) and 67 other genes trended upward.

Remarkably, of the 777 genes we had assigned to various expression modules using mouse myeloid datasets^8^, only five were detected as differentially expressed in SFG myeloid cells from AD patients: *PLXNC1*, *TGFBI*, and *ADAM8*, along with *APOE* and *SERPINF1*. In addition to *ADAM8*, two other genes from the LPS-related gene set showed strong upward trends (*TSPO*, P = 0.14 and *CD44*, P = 0.11), while *PILRA* (P = 0.63) and 58 other genes from this set showed only weak upward (35) or downward (24) trends. All of the DAM, Brain Myeloid, and LPS-related genes named above except *APOC1* and *TGFBI* were included in our qPCR panel, and these genes’ RNA-Seq measurements from SFG myeloid cells were generally reproduced in FuG myeloid cells except for *ADAM8* (Extended Data Fig. 3b).

We next asked whether the human AD genes we identified (besides *APOE*, a well-known DAM gene) showed consistent trends in mouse models of neurodegeneration^8–10^, infection^6,11^, and aging^12^. Of the Myeloid-AD-Up genes, *PLXNC1*, *CD44*, *SMIM3*, and *ADAM8* were frequently though modestly increased in neurodegenerative mouse models (Extended Data Fig. 6). Of the Myeloid-AD-Down genes, only *SERPINF1* showed consistent reduction in these models. Due to the general lack of overlap with the DAM activation state in mouse AD models, or with any known activation states in any models, we refer to unique combination of upregulated and downregulated genes we identified in <u>h</u>uman <u>A</u>D <u>m</u>icroglia/myeloid cells as the HAM profile.

We next examined human microglia expression profiles published by other groups for possible relationships with the HAM signature. We found interesting correlations between our dataset and the microglia profiles obtained from fresh autopsy tissue by Galatro and co-workers^1^. Although our subjects tended to be mostly older (Extended Data Fig. 7a), we confirmed Galatro’s age-associated DE in our own myeloid cell profiles from control subjects (Fig. 3b). Strikingly, we also found a relationship between age-associated DE and AD-associated DE. Genes higher in brain myeloid cells from older subjects, like *IL15*, tended to be elevated in AD relative to controls, and genes lower in cells from older subjects, like *CECR2*, tended to be reduced in AD relative to controls (Fig. 3c, d). This was not due to differences in age between our AD and control subjects (Extended Data Fig. 7a). On the other hand, roughly half of the DE genes in our dataset, including *APOE* and *LSR*, showed no relationship with age in microglia. This suggests that the HAM profile in AD microglia might reflect a mixture of an “enhanced aging” process as well as an age-independent, disease-related activation process.

We also analyzed another human dataset, from Gosselin and co-workers^2^, of microglia expression profiles obtained from fresh surgical tissue (all from young subjects, see Extended Data Fig. 7a) and blood monocytes from the same subjects. DE genes between monocytes and microglia correlated reasonably well between human and mouse^13^ datasets (Extended Data Fig. 7b). Comparing the two human datasets, we saw that many of the genes elevated in myeloid cells from younger subjects in Galatro’s dataset were also microglia-enriched in Gosselin’s dataset, like *CECR2*; conversely, many genes elevated in myeloid cells from older subjects were monocyte-enriched, like *IL15* (Fig. 3d, Extended Data Fig. 7c). This may suggest that some of the age-related changes could be due to increased brain infiltration of peripheral monocyte-derived cells. Alternatively, these changes could simply reflect microglial transcriptional modulation toward a state that bears a slight resemblance to monocyte profiles. Putting all three datasets together, we can categorize our HAM signature genes into modules according to whether they increase or decrease in AD, whether they vary with age, and whether they are microglia- or monocyte-enriched (Extended Data Fig. 7d, Supplementary Table 2). These modules may represent different biological processes.

Finally, we analyzed a recent dataset of single cell RNAseq profiles obtained from tumefactive MS lesions or healthy control tissues for evidence of DAM and HAM profiles in another human disease setting^14^. From each cell cluster, we aggregated cells from the same subject and batch into a single expression profile (Extended Data Fig. 8a), and then analyzed these sample populations for expression of mouse DAM genes, our Myeloid-AD-Down genes, and our Myeloid-AD-Up genes. Roughly half of the DAM genes showed elevated expression in the cells corresponding to one or more of the authors’ activated microglia clusters Hu-C2, Hu-C3, or Hu-C8 (Extended Data Fig. 8b). This evidence that human microglia are capable of a DAM-like response makes its absence in our AD dataset more notable. More than half of our Myeloid-AD-Down genes appeared reduced in microglia from MS lesions (Extended Data Fig. 8c), suggesting that their downregulation might be a common feature in different disease settings. About one-third of the Myeloid-AD-Up genes were expressed more highly in at least one of the activated microglia clusters compared to control microglia (Extended Data Fig. 8d), suggesting that several of the genes we identified as upregulated in AD myeloid cells, including the transcription factors *FOXP1* and *RUNX3*, are also employed in other neurodegenerative disease contexts. In summary, microglia from tumefactive MS lesions showed partial overlap with both the DAM and HAM profiles.

Here we have described the AD-associated DE profile of the brain myeloid compartment. We cannot exclude that our experimental methods somehow precluded the detection of human microglia with DAM-like activation, but the stronger HAM signal observed in whole tissue RNA underscores the relevance of our findings. Despite the dissimilarity between DAM and HAM signatures, some qualitative similarities emerge. First, just as the DAM genes induced in neurodegenerative mouse models overlap with those induced by natural aging^8,15^, so do many of the HAM genes induced in human AD tissues (Fig. 3b), though the genes involved are completely distinct between species^1^. Second, an emerging theme in recent mouse DAM literature is the role of lipid and lysosomal biology: many mouse DAM genes such as *apoe*, *Ch25h*, *Lpl*, *Ctsb* and *Atp6v0d2* are known to function in lipid and lysosomal biology and can be induced by lipid pathologies including demyelination^16^ and atherosclerosis^17,18^. In our data, in addition to *APOE*, we found that the lipoprotein receptor *LSR* and the lysosomal enzyme *ARSA*—homozygous mutations in which cause metachromatic leukodystrophy^19^—are elevated in myeloid cells from AD patients. Therefore, another qualitative commonality between DAM and HAM genes could be the involvement of lipid/lysosomal biology-associated genes. Several genes associated with AD incidence (*APOE*, *CLU*, *ABCA7*, *SORL1*, *INPP5D*, *PLCG2*) also function in lipid transport and signaling^20–22^.

Why are the HAM and DAM gene signatures so different? One explanation could be intrinsic differences in human versus mouse innate immune responses, but the activation of many DAM genes in MS lesions suggests this is not the main reason. Another explanation could be the very different stages of disease being analyzed, with mouse β-amyloid models perhaps representing early stage AD with amyloid deposits present but preceding neurodegeneration. However, if this were the main reason, we might expect to see mouse DAM genes elevated in early Braak stage tissues and decreased in later Braak stages, but we have not observed such trends in whole tissue RNA profiles (Extended Data Fig. 4c). A third explanation for the dissimilarity could be that the DAM activation state in β-amyloid models is a protective response by healthy microglia^23^, whereas genetic and histological findings suggest that human AD involves impairments in microglial activation^24,25^. Additional profiles with increased cellular resolution for various AD stages and brain regions, different neurodegenerative diseases, and new disease models that incorporate human microglial cells will shed further light on how the HAM profile relates to mechanisms of AD protection or pathogenesis.

## Supporting information

Supp Table 1: Sample-Level Quantification

Supp Table 2: Group-Level Quantification

---

**Extended Data Figure 1.**
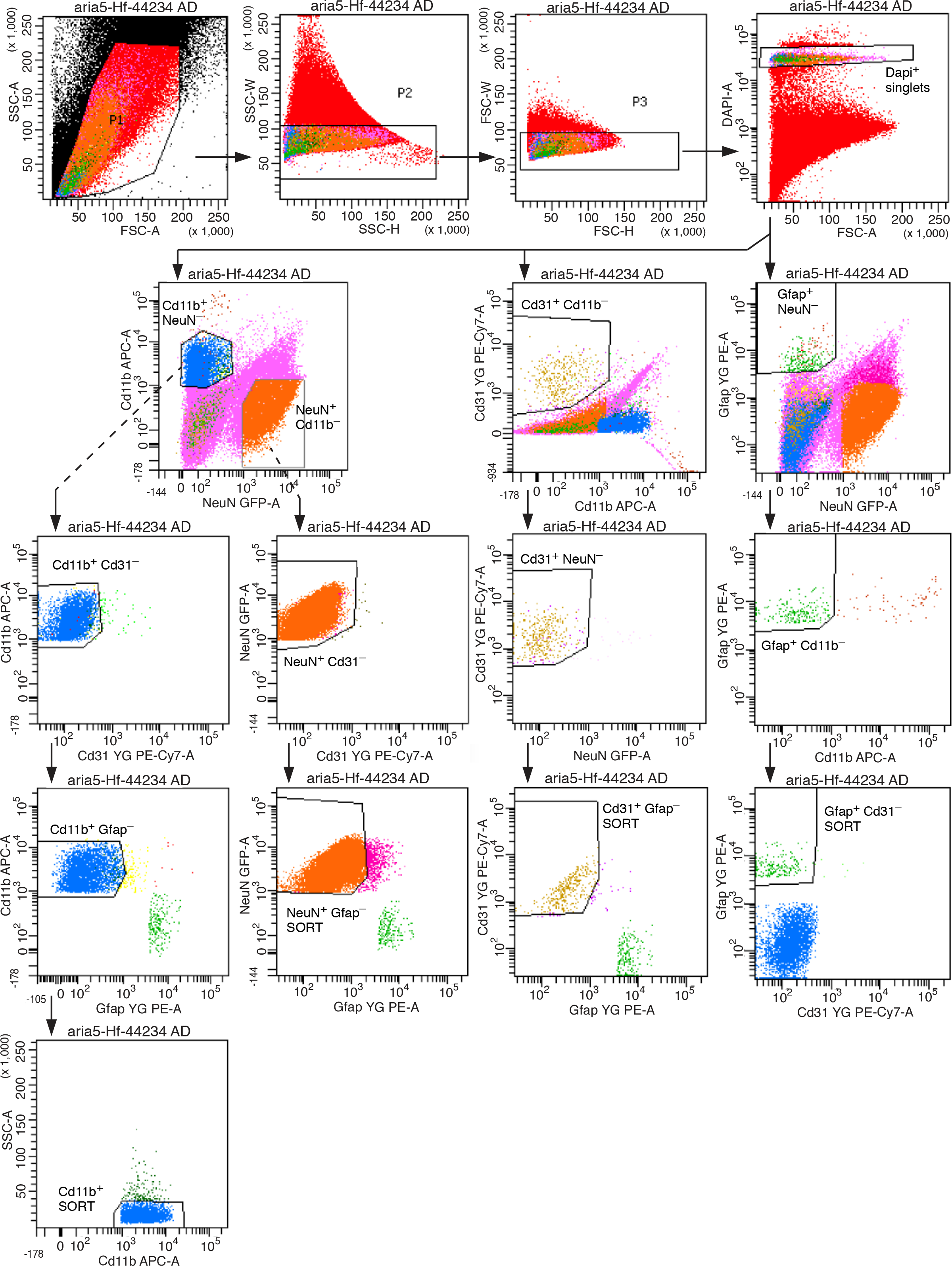
Example FACS plots showing isolation of four cell type populations from one AD sample.

**Extended Data Figure 2.**
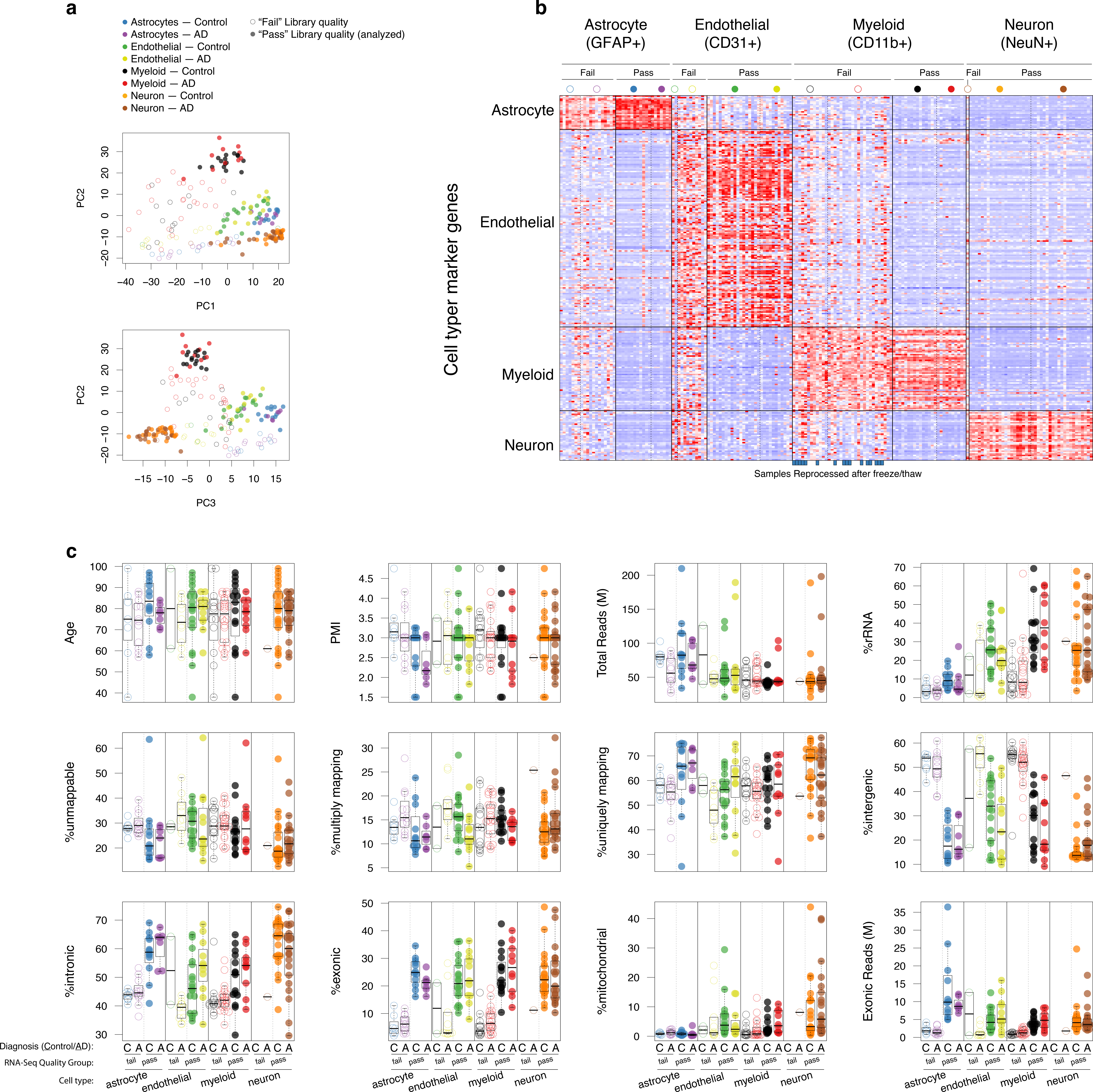
Sample quality control. (a) PCA analysis of passing and failing RNA-Seq profiles, corresponding to Fig. 1c. Note that separation of cell types degrades in “Fail” samples. (b) Heatmap of all samples. Solid vertical lines separate cell types, and Pass/Fail within cell types. Dashed lines separate Control and AD samples. Libraries from 16 samples which were repeated after freeze/thaw are indicated below. (c) Subject, sample and library attributes separated by cell type, quality group (Pass/Fail) and diagnosis. Age and PMI in particular do not appear to show strong differences between Pass and Fail samples, although several RNA-Seq library statistics, such as %intergenic, do appear quite different. Denominators for percentages as follows: %rRNA, total reads; %unmappable, %multiply mapping, %uniquely mapping, “processed” reads (total reads with rRNA, low quality, and adapter contamination removed); %intergenic, %intronic, %exonic, %mitochondrial: total uniquely mapping reads.

**Extended Data Figure 3.**
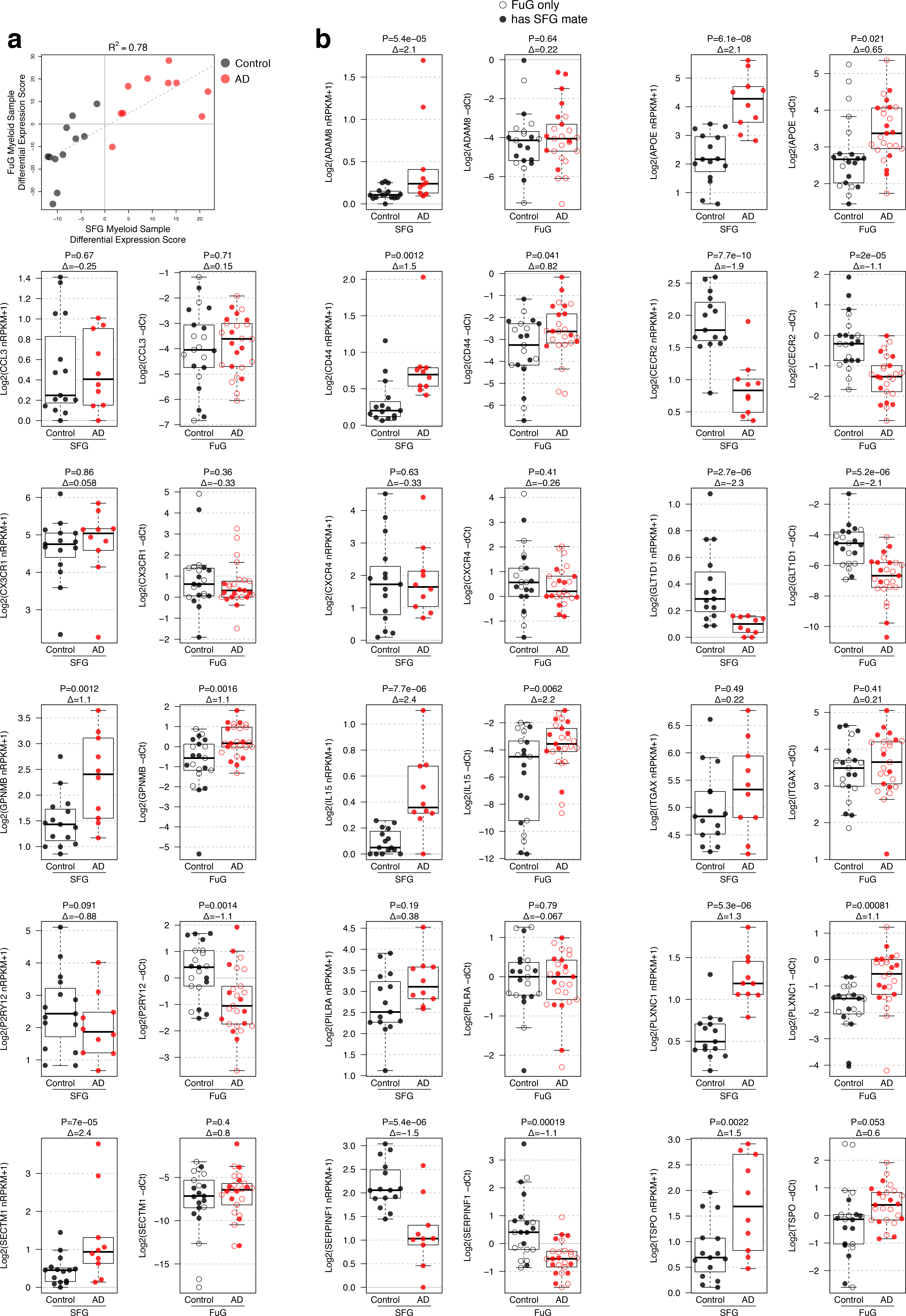
AD brain myeloid differentially expressed (DE) genes from SFG RNA-Seq data are largely reproduced in FuG qPCR data. (a) DE scores were calculated for sorted myeloid cells from SFG and FuG of the same subjects, with higher scores indicating increased degree of differential expression for the genes identified as DE by RNA-Seq in the SFG samples and present in the qPCR panel. Each point represents one subject for which passing SFG RNA-Seq and FuG qPCR profiles were available, with coordinates giving the DE scores (Methods) for corresponding SFG and FuG profiles. (b) Selected examples of gene expression measurements in SFG myeloid cells by RNA-Seq and FuG myeloid cells by qPCR.

**Extended Data Figure 4.**
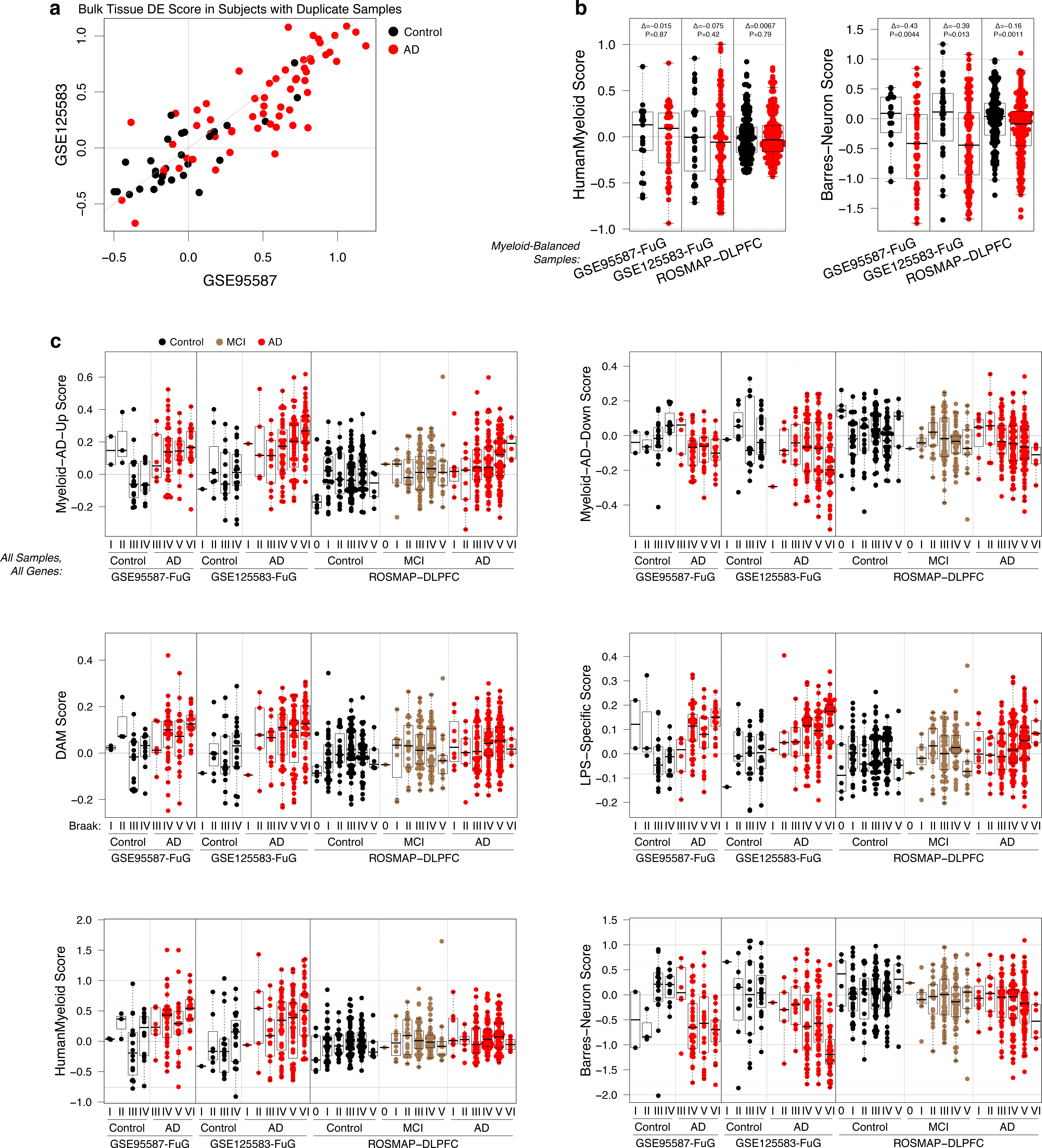
Bulk tissue analyses. (a) Duplicated samples show consistent DE. 89 samples were duplicated in GSE95587 and GSE125583, in the sense that they came from different tissue blocks of the same fusiform gyrus. For each of these, a samplewise DE score was calculated separately in the two datasets using common bulk DE genes. Plot shows that the DE scores are highly correlated, indicating that the expression signature of a small piece of tissue reflects the entire brain region. (b) Myeloid-balancing results in similar distributions of myeloid scores but still a strong depletion of neuron gene expression in bulk AD brain tissue. Plot shows gene set scores of indicated gene sets in individual bulk tissue samples from three different cohorts, similar to Fig. 2d but for different gene sets. Also, neuronal genes were not removed (that would be pointless in this context since none of the HumanMyeloid and all of the Barres-Human genes are neuronal). (c) Human AD Myeloid gene changes are observed in bulk tissue at later Braak stages. Plots are similar to Fig. 2d but include all samples and genes, without myeloid balancing or removing neuronal genes, with samples stratified by Braak stage.

**Extended Data Figure 5.**
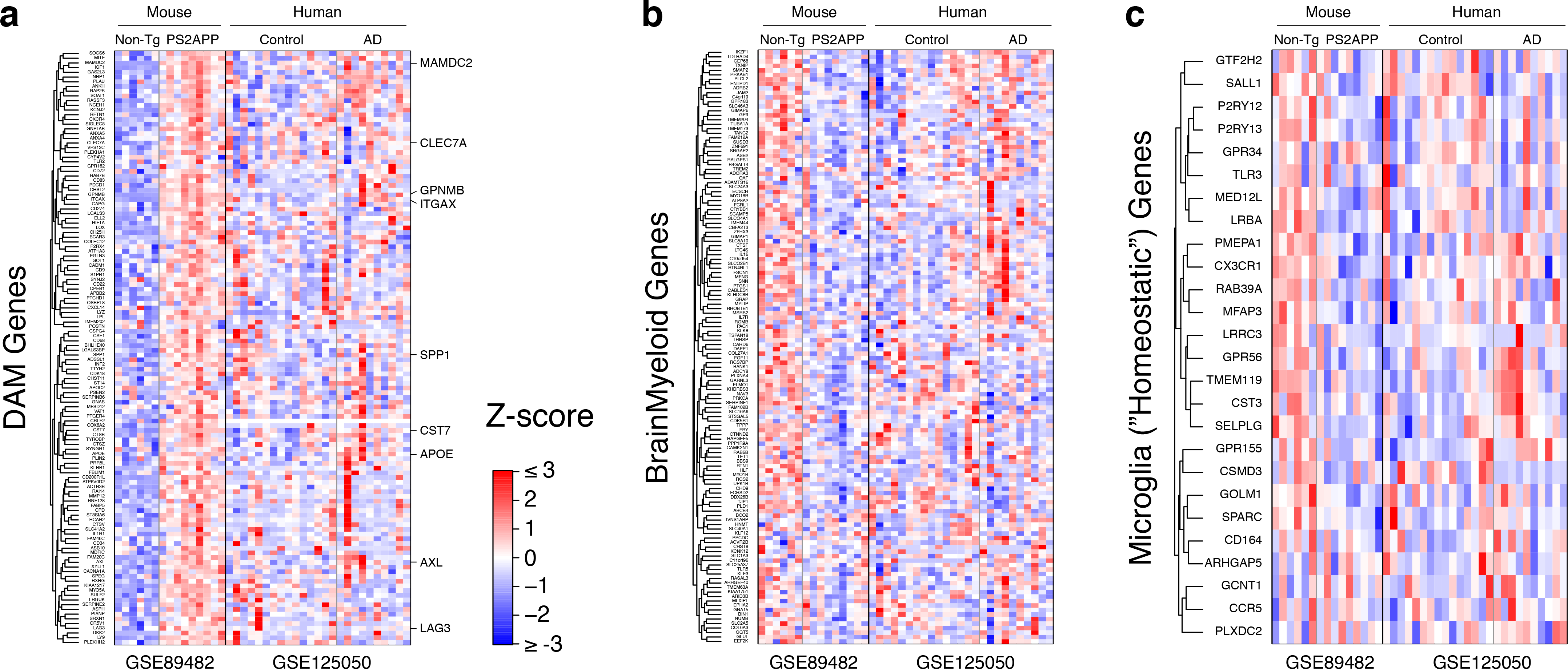
Heatmaps of (a) DAM, (b) Brain Myeloid, (c) homeostatic Microglia gene set in brain myeloid cells from the PS2APP mouse model of β-amyloidosis (GSE89482) and human SFG myeloid cells from this study (GSE125050).

**Extended Data Figure 6.**
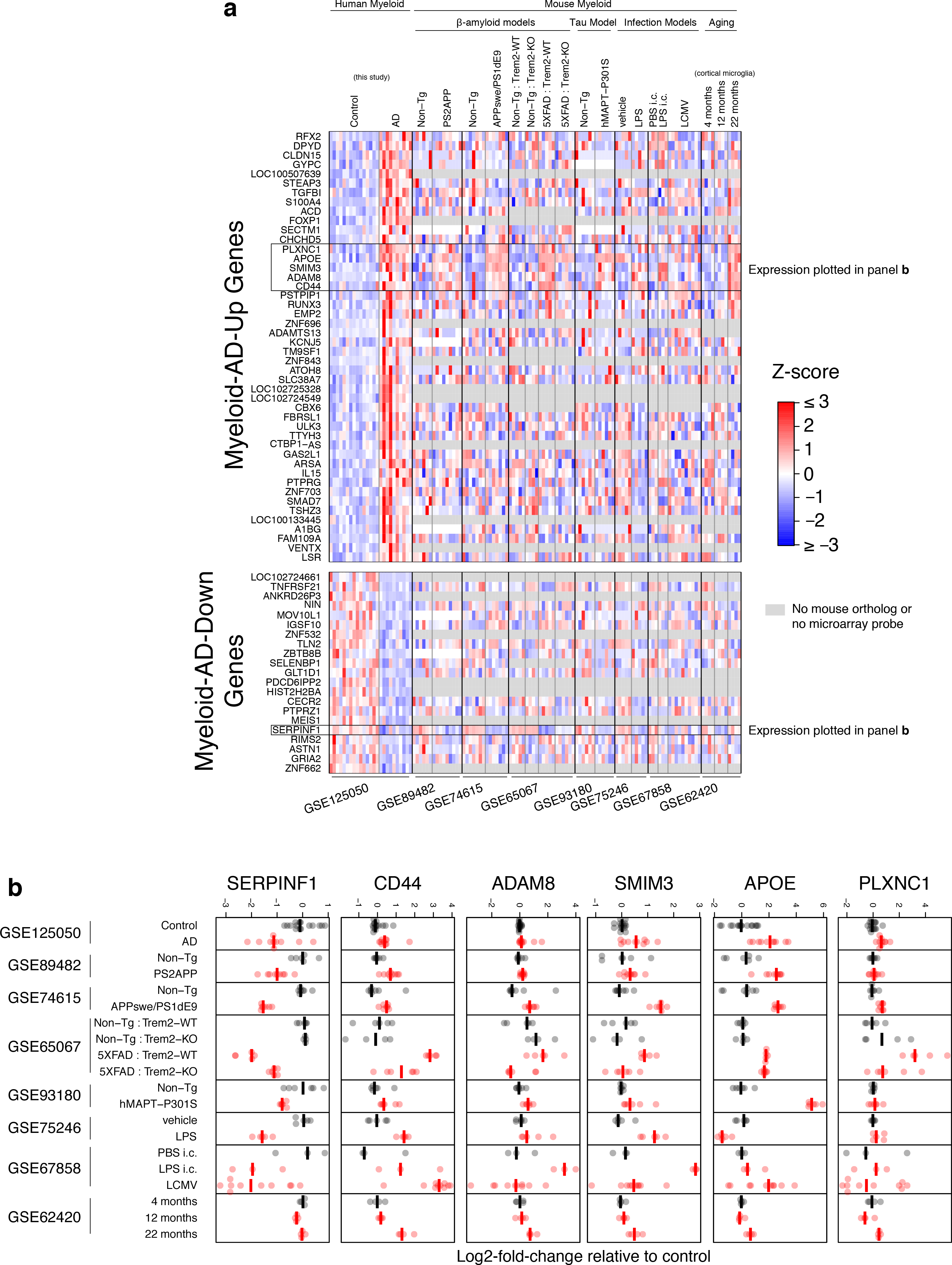
(a) Heatmaps of human AD myeloid DE genes in mouse myeloid data sets. (b) Expression of selected genes (the few which exhibit consistent changes in mouse data sets) in individual samples in mouse data sets. Expression values are normalized to average expression in the control group within each study.

**Extended Data Figure 7.**
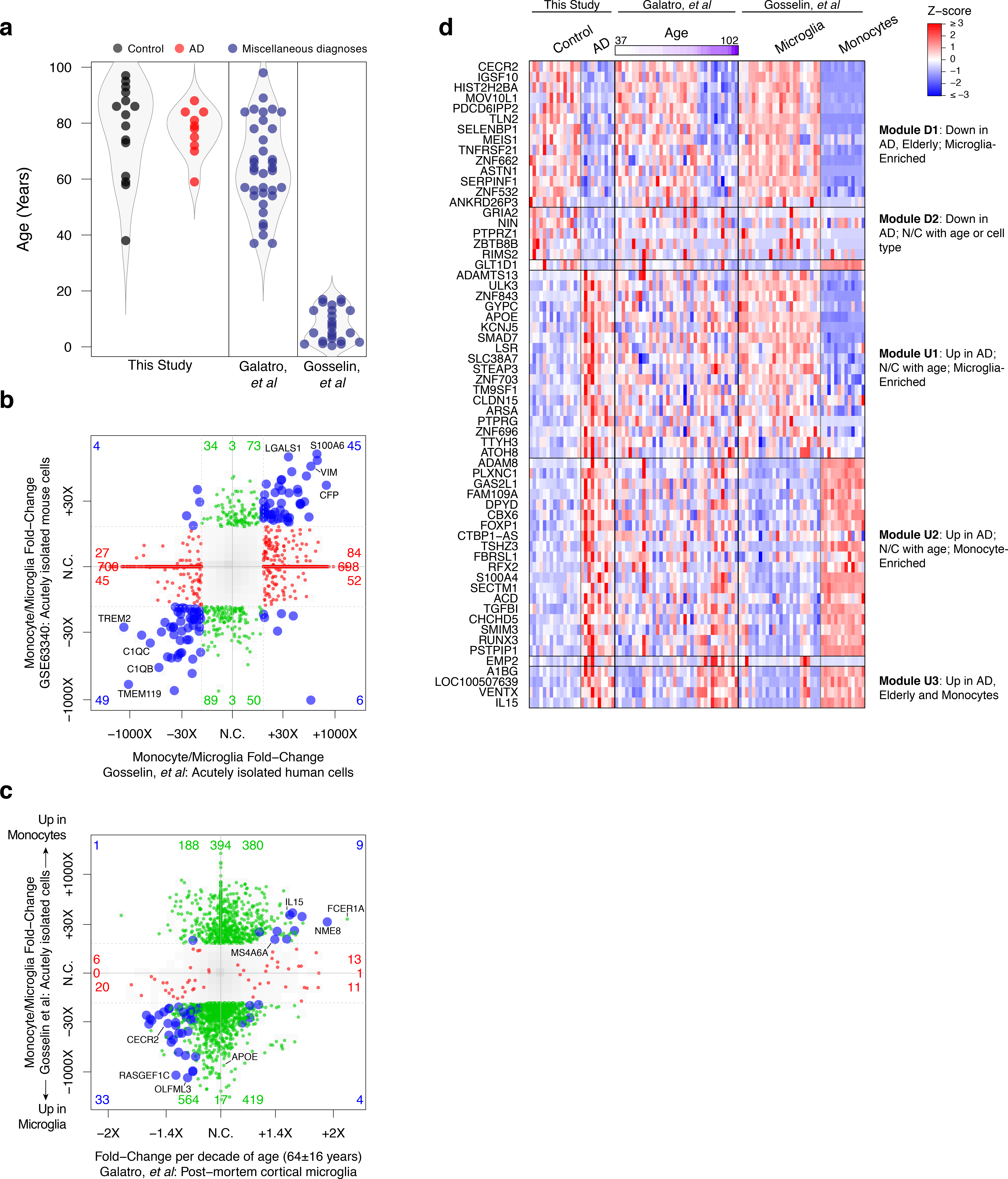
Monocyte/Microglia DE genes may contribute to both late aging and AD signatures. (a) Distribution of subject ages in three studies. (b) Monocyte/microglia DE profile is similar in human ^2^ and mouse ^13^ studies. “4-way” plot, like Fig. 3a, but DE between monocyte and brain myeloid (“microglia”) profiles is shown. (c) Many DE changes elevated or depleted with aging (x-axis, red and blue genes to the right or left, respectively) are also elevated or depleted, respectively, in blood-derived monocytes relative to microglia (*y*-axis). (d) Heatmap and clustering of AD DE genes in three datasets. Gene modules and order were defined based on DE and direction in each of the three datasets, as indicated. *GLT1D1* and *EMP2* are not contained within any module. Modules are indicated in Supplementary Table 2.

**Extended Data Figure 8.**
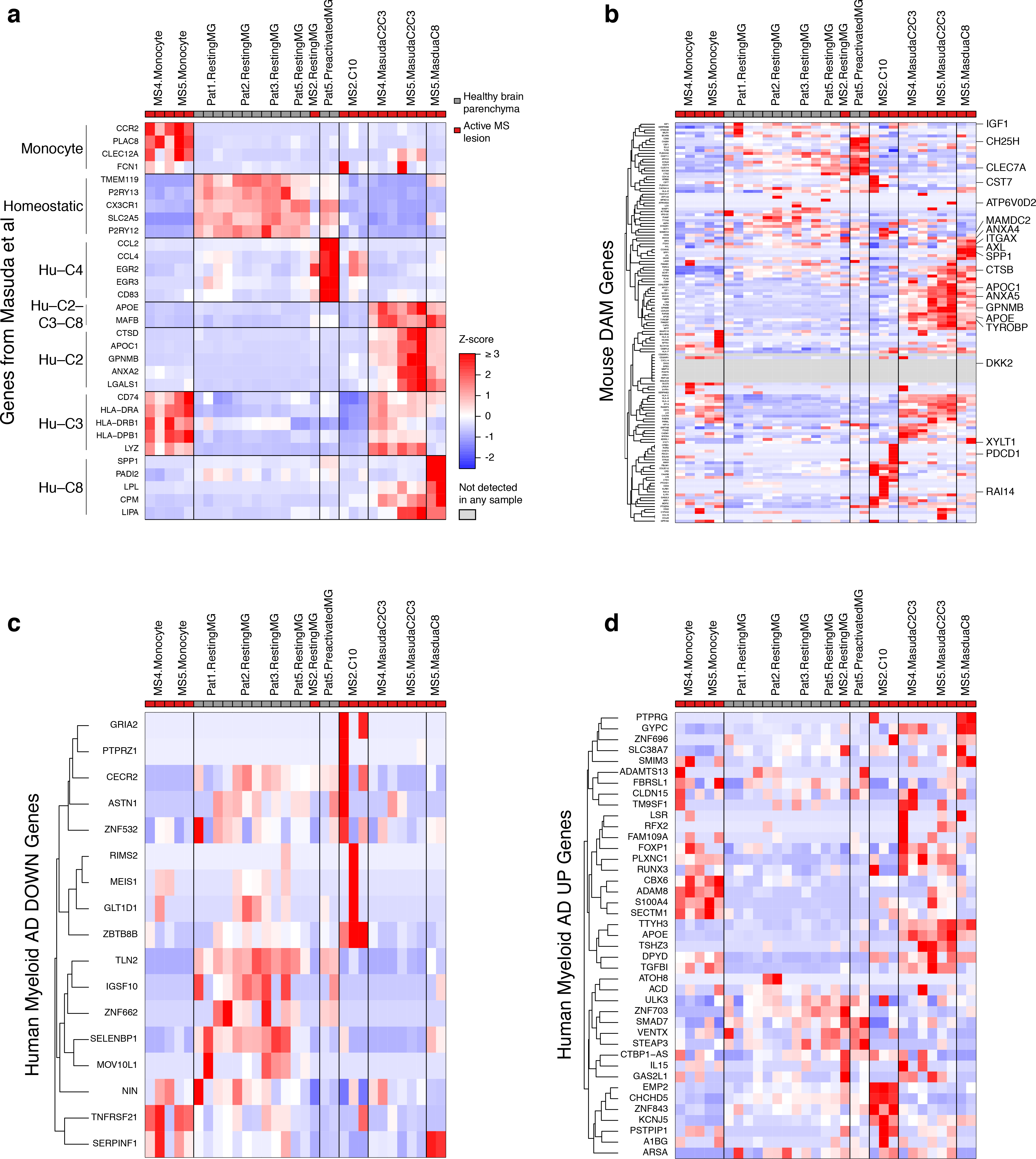
Pseudobulk analysis of human brain myeloid single-cell datasets from Masuda *et al* ^14^. We ran clustering analysis using Seurat and used gene sets listed in the other paper to associate our clusters with the authors’ clusters. We then created “pseudobulk” profiles summarizing each interpreted cell type within each sequencing multiplex. The four panels show heatmaps of gene expression (Z-score of normalized count) of different gene sets in each pseudobulk. Pat1, Pat2, Pat3, Pat5 are subject identifiers for the normal tissue samples and MS2, MS4 and MS5 are for the three MS lesions. C10 is a cluster arising from our analysis that we could not unambiguously assign to Masuda’s clusters, but which consisted mostly of cells from subject MS2. MasudaC2C3 is a cluster arising from our own analysis that seemed to consist of cells from clusters C2 and C3 as named by Masuda, and MasudaC8 is the cluster from our analysis that clearly corresponded to the cluster C8 from Masuda. (a) Gene sets listed in Masuda *et al*, to identify the clusters from our analysis. (b-d) DAM, AD-Down and AD-Up gene sets. Selected genes are re-labeled on the right in (b) for clarity.

Supplementary Table 1. Sample-level gene expression values. Excel file (52 MB) with one row per gene. Columns give normalized sample-level gene expression values and differential expression statistics for GSE125050 (SFG RNA-Seq from this study), the fusiform gyrus qPCR (also from this study, but not deposited into GEO) as well as the other human sorted cell datasets analyzed, (phs001373.v1.p1 “Gosselin”, GSE99074 (“Galatro”) and GSE73721 (“Barres”).

Supplementary Table 2. Gene annotations, group-level gene expression and genome-wide DE results. Excel file (24 MB) with one row per gene. Columns give membership in all the gene sets described in this manuscript, average gene expression values by group, as well as differential expression statistics for the studies in Supplementary Table 1 as well as mouse datasets GSE75431 (sorted cells from PS2APP mouse), GSE93180 (sorted microglia from Tau-P301S mouse), GSE75246 (sorted microglia from LPS-treated mice), GSE95587+GSE125583 (two cohorts of bulk AD patient FuG) and ROSMAP-DLPFC (bulk AD patient DLPFC).

## Methods

### Human Patient Tissue Samples

Frozen superior frontal gyrus and fusiform gyrus tissue blocks and pathology/clinical reports, including age, sex, diagnosis, and Braak stage, were obtained from the Banner Sun Health Research Institute Brain and Body Donation Program in accordance with institutional review boards and policies at both Genentech and Banner Sun Health Research Institute.

All subjects were characterized clinically and neuropathologically by the Arizona Study of Aging and Neurodegenerative Disease/Brain and Body Donation Program ^26^. All Alzheimer’s disease subjects were clinically diagnosed with Alzheimer’s disease in life and brains were neuropathologically confirmed to have “frequent” CERAD neuritic plaque densities ^27^ and Braak score V or VI ^28^. Controls did not have dementia, AD or other neurological disease diagnoses in life.

For sorted cell cohort (GSE125050), controls had either “zero” or “sparse” CERAD neuritic plaque densities, and had Braak scores ranging from 0 to IV (median II). One control subject of Braak stage II was diagnosed post-mortem with “argyrophilic grain disease”.

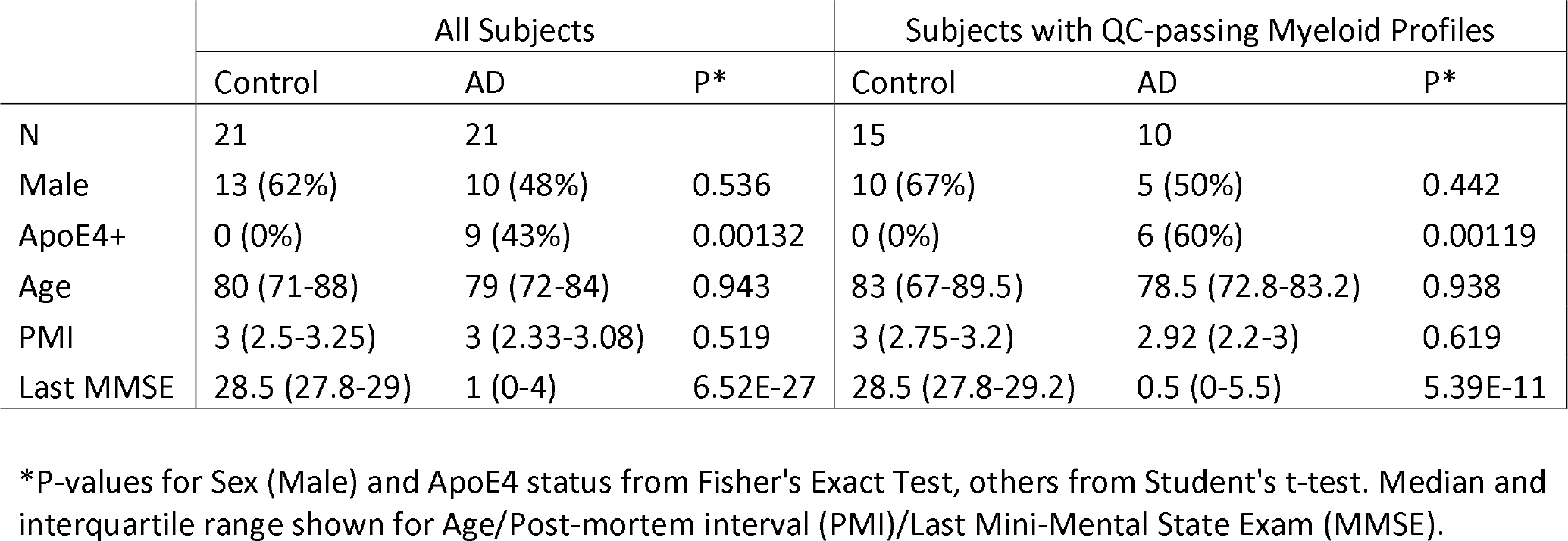

Visual exploration as well as linear model testing reveal no significant correlation between PMI and any of the other variables (sex, *APOE*4 status, age, Last MMSE, or diagnosis).

Bulk tissue studies cohorts were as follows:

|  | GSE95587 (previously published) |  |  | GSE125583 (new subjects in this study) |  |  |
| --- | --- | --- | --- | --- | --- | --- |
|  | Control | AD | P* | Control | AD | P* |
| N | 33 | 84 |  | 42 | 158 |  |
| Male | 23 (70%) | 42 (50%) | 0.0644 | 19 (45%) | 86 (55%) | 0.3 |
| ApoE4+ | 8 (24%) | 38 (45%) | 0.0574 | 5 (12%) | 85 (54%) | 5.29E-07 |
| Age | 82 (80-90) | 87 (81-91) | 0.471 | 89 (84.2-91) | 84 (77-88) | 7.45E-06 |
| Last MMSE | 29 (28-29) | 17 (7-22) | 2.10E-23 | 28.5 (27-30) | 14 (6-21) | 1.06E-47 |

\*P-values for Sex (Male) and ApoE4 status from Fisher's Exact Test, others from Student's t-test. Median and interquartile range shown for Age/PMI/Last MMSE.

### Tissue Processing, Library Preparation, and RNA-Seq

All samples obtained from Banner Research were stored at −80ºC until the time of processing. For bulk tissue studies (GSE125583), frozen tissue was sectioned in approximately 8 slices 40 μm thick and stored at −80ºC. Tissue was homogenized in 1 ml QIAzol with 5 mm stainless steel beads using a Tissuelyzer (20 Hz for 4 min). After homogenization, 200 μl of choloroform were added to the cleared lysate (1 min 12,000 rcf at 4ºC), vigorously shook and incubated at room temperature 2-3 min. Samples were centrifuged for 15 min at 12,000 rcf at 4ºC and the upper aqueous phase was transferred to a new tube. RNA was extracted using Qiagen miRNeasy mini columns, yielding samples with RNA integrity (RIN) scores averaging 6.5. Standard polyA-selected Illumina RNA-seq analysis was performed as described ^6^ on samples with RNA integrity (RIN) scores at least 5 and post-mortem intervals (PMIs) no greater than 5 hr. Of 289 total samples, 89 were from subjects that had already been profiled in our previous study, GSE95587. These are available in GSE125583 and marked therein as duplicated in GSE95587. These samples, which came from new fusiform gyrus tissue blocks, showed very similar sample-by-sample DE profiles as the corresponding samples from the same subjects in GSE95587 (Extended Data Fig. 4a), but were omitted in all other analyses associated with this manuscript to avoid overlap between the two datasets (Fig. 2d, Extended Data Fig. 4b-c, supplementary tables and website).

For sorted cell studies, frozen samples were opened on dry ice and a 100-200 mg portion was excised. The excised portion was thawed in ice-cold Hibernate A and minced on a cold block with a pre-chilled razor. The tissue was then transferred to a 2 ml round-bottom tube with cold 1.6 ml of Accutase. Minced SFG samples included both gray and white matter, while only gray matter from FuG was used for mincing. (For sixteen SFG samples, excess minced tissue fragments were refrozen and stored for a later attempt to repeat the entire sorting and RNA-Seq procedure from the same brain region—see QC section below.) Minced samples were incubated 20-30 minutes on a rotator at 4ºC, mechanically dissociated, centrifuged and resuspended, and ethanol-fixed for 10 minutes on ice as previously described ^6^. Cells were washed briefly and incubated with anti-CD11b APC (Millipore MABF366), anti-GFAP PE (BD Pharmingen 561483), anti-NeuN AlexaFluor488 (Millipore MAB377X), anti-CD31 PE-Cy7 (BD Pharmingen 563651), and Human Fc Block (BD Pharmingen 564220) for 20 minutes at 4ºC with sample rotation. Cells were centrifuged at 2,000 rcf for 2 minutes and briefly washed prior to DAPI (1 mg/ml stock) being added at 1:1,000 followed by FACS sorting on ARIA sorters. Only DAPI+ singlet cell bodies were collected, and each cell population of interest was gated to be negative for all the other antibody marker channels. Samples were generally processed in pairs, with one AD and one control sample. While each human sample was unique and gating was occasionally fine-tuned, samples generally separated based on the same broad FACS gates. Typical cell numbers collected were 100K CD11b+ cells, 40K GFAP+ cells, 10K CD31+ cells, and 400K NeuN+ cells. FACS-isolated cell populations were spun at 5,000 rcf for 5 minutes and resuspended in 0.35 ml Buffer RLT from Qiagen RNeasy Micro kit. Lysed samples were stored at −80ºC until all samples for a given brain region were sorted. Each cell type was then processed as a single batch. Total RNA extracted from sorted cell populations was subjected to Fluidigm qPCR assay which yielded reliable cell-specific gene expression data, despite subpar RNA quality resulting from post-mortem status, freeze/thaw process and fixation. In addition to the methods for dissociation and immunolabeling described above, we also attempted dissociation techniques involving trypsin or papain at 37ºC, psychrophilic proteases at 4ºC, longer Accutase treatment periods, automated mechanical dissociation instead of pipetting, other fixatives besides ethanol, labeling and sorting of non-fixed cells for cell types with surface markers (CD11b and CD31), and antibodies for several alternative cell type markers. None of these attempts were as good as the method described above in terms of cell yield and RNA recovery. RNA integrity (RIN) and concentration was determined by 2100 Bioanalyzer (Agilent Technologies). RIN scores for all cell types were typically between 1 and 3. Typical RNA yields were 1 µg for neurons, 25 ng for microglia and astrocytes, and 5 ng for endothelial cells. cDNA was generated using Nugen’s RNA-seq method for low-input RNA samples, Ovation RNA-seq System V2 (NuGEN). We chose the Nugen kit to enable recovery of 5’ mRNA sequences since it uses random oligos for cDNA priming, given the highly fragmented condition of our sorted cell RNA preps. Generated cDNA was sheared to 150-200bp size using LE220 ultrasonicator (Covaris). Following shearing, the size of cDNA was determined by Bioanalyzer DNA 1000 Kit (Agilent) and quantity was determined by Qubit dsDNA BR Assay (Life Technologies). Sheared cDNA was subjected to library generation, starting at end repair step, using Illumina’s TruSeq RNA Sample Preparation Kit v2 (Illumina). Size of the libraries was confirmed using 4200 TapeStation and High Sensitivity D1K screen tape (Agilent Technologies) and their concentration was determined by qPCR based method using Library quantification kit (KAPA). The libraries were multiplexed within cell types and then sequenced on Illumina HiSeq2500 (Illumina) to generate 50M of single end 50bp reads.

### RNA-Seq Data Processing and QC

Sorted cell and bulk RNA-Seq data were analyzed as described ^8^, except as follows. For Gosselin *et al* we did not have access to the raw FASTQ files, so we used the author-provided tables of counts and TPM values. For ROSMAP-DLPFC we downloaded the file ROSMAP_RNAseq_FPKM_gene_plates_1_to_6_normalized.tsv from the synapse website, in order to take advantage of the batch normalization that the authors already applied. We did not used the samples from batches 7 and 8 since, despite restricting to the batch-normalized values, we still saw very strong clustering of these two batches separately from the first 6 on PCA. For Masuda et al, we downloaded each of the 32 gene quantification files from the GSE124335 GEO record (file names like GSM3529822_MS_case1_3.coutt.csv.gz). These files each contained 192 columns corresponding to the cells of one batch, and one row per gene. The gene labels were mapped onto our internal gene annotation (based on Ensembl) first by trying to match symbols and then aliases. After this step cells with less than 800 total transcripts or greater than 30% mitochondrial transcripts were discarded, resulting in 1738 QC-passing cells for analysis. Total transcript number normalization was performed, dividing each gene expression value for a cell by a factor proportional to the total number of transcripts in that cell.

“Pass” or “Fail” status for our sorted cell RNA-Seq profiles was determined by a combination of PCA analysis and heat map analysis (similar to the heat map in Extended Data Fig. 2b but with unbiased hierarchical clustering). Failed libraries generally showed much higher percentages of intergenic reads, and lower percentages of exonic and intronic reads. Library failure was not random: when one cell type library from a tissue failed, other cell type libraries from the same tissue were more likely to also fail, despite the fact that RNA purification, library prep, and sequencing were performed at different times for each cell type. In addition to pre-existing conditions of frozen tissues, other factors involving low RNA integrity and quantity contributed to the likelihood of library failure. RNA integrity was negatively impacted both by whole tissue freeze/thaw and EtOH fixation of dissociated cells. Samples giving rise to failed libraries passed the initial qPCR screening for cell type markers since the ΔCt scores for markers relative to housekeeping genes were normal (although raw Ct values were of course higher in low quantity samples); therefore, aspects of the cDNA synthesis and amplification procedure also contributed to library failure. Most neuron libraries were “Pass” since the higher number of cells and quantity of RNA collected was enough to overcome the limitations of low RNA integrity. The challenges introduce by using frozen tissues became evident when we processed the sixteen refrozen portions of minced SFG tissue to test whether we could get reproducible RNA-Seq data from tissues frozen multiple times. Unfortunately, 100% of sorted cell libraries prepared from refrozen tissues failed (despite passing initial QC by cell type marker qPCR). This suggested that factors related to freezing and sampling of the initial tissues prior to their arrival at Genentech may have contributed to the rate of library failure in the first round of RNA-Seq.

Principal Components Analysis (Fig. 1, Extended Data Fig. 2a) was performed on Z-score normalized matrix of 1000 most variable genes by IQR using the R function `prcomp()`.

### Differential Expression (DE) Analysis

DE between AD and controls for this study’s sorted cell populations was first attempted using voom+limma, which identified only 12 DE genes (adjusted P ≤ 0.05) in myeloid cells and none in the other cell types. We then tried using DESeq2 (adjusted P ≤ 0.05, with maximum Cook’s *P*-value ≥ 0.01 filter to help exclude genes driven by outlier samples), which identified greater numbers of DE genes (4 in neurons, 66 in myeloid cells, and 135 in endothelial cells). In the myeloid cells, 11/12 DE genes identified by voom+limma were also identified by DESeq2, with *CD44* being the only exception (P=0.113 in DESeq2). We included *CD44* in our panel of genes tested by qPCR in FuG myeloid cell sorts, and it was again increased in the AD samples (P=0.036), so we consider its DE to be genuine. As for the larger numbers of DE genes identified by DESeq2, the absence of any voom+limma hits for neurons and endothelial cells and the lack of other human AD datasets suitable for cross-comparison led us to set these cell types aside for now (taking a conservative position) and focus on the changes in myeloid cells which were corroborated by FuG qPCR, overlap with the human aging dataset, and whole tissue RNA profiles.

Our analysis of Galatro *et al* was performed using DESeq2, filtering results for maximum Cook’s *P*-value ≥ 0.01 to exclude outlier-driven hits. For Galatro *et al* the ages of the subjects were taken from their supplemental table rather than GEO (these differed only for the sample GSM2631906), and the DE analysis was simply the linear model `~Age`, only using the samples with `tissue=“Microglia”`. For Gosselin et al, DE between microglia and monocytes was performed using `voom+limma` only using the samples with `CultureStatus=“ExVivo”`.

### Other Covariates

Linear modeling revealed that the expression of about 80 genes was significantly increased in our human myeloid cells from subjects with larger post-mortem interval (PMI; data not shown). This seemed to be largely driven by elevated mitochondrial gene expression in a subset of the samples with large PMI. However, the distribution of PMI in our AD and control samples was similar (Extended Data Fig. 2c), there was no overlap between the DE genes and the PMI-associated genes, and adding PMI to our statistical model for AD-associated DE gave very similar results. Therefore, we did not include PMI in subsequent analyses. We only detected one DE gene, *ACY3*, between the *APOE*4- and *APOE*4+ AD myeloid cells. It showed variable expression levels in the Controls, so it may be a false positive. Sex-associated DE in myeloid cells was only seen for a few X and several Y chromosome genes.

### Fluidigm qPCR Analysis

qPCR data were collected as described ^6^. Then, for each assay target, the maximum *C*_*t*_ of quality > 0 was calculated. The *C*_*t*_ value maxCt+0.5 was assigned to each assay that had *C*_*t*_ larger than this value (including 999). All assays were performed in duplicate and the average of these two *C*_*t*_ values was kept, except for twelve sample/assay pairs for which the difference was more than 2.82 (corresponding to a standard deviation of 2), which were discarded. *ΔC*_*t*_ normalization was performed using global median (the median Ct value for all assays for a given sample) and differential expression between AD and control was performed using limma.

### Gene Set Analysis

Cell type marker gene sets in Fig. 1b were previously described ^8^, available in column N of that manuscript’s Supplementary Dataset S5, and repeated in column O of this manuscript’s Supplementary Table 2.

Most of the gene sets used in Figs. 2d, 3a, and Extended Data Figs. 4, 5, 6, and 8 were previously described ^8^, available in columns U and V of that manuscript’s Supplementary Dataset S4, and repeated in this manuscript’s Supplementary Table 2, but summarized briefly here, along with any relevant differences from the previous manuscript:

- Interferon-Related: interferon-stimulated genes. Example: *IFIT1*.
- DAM: disease/damage-associated microglia genes (called “Neurodegeneration-Related” in the previous manuscript). Example: *CST7*.
- Microglia: microglia-specific genes, elevated in microglia relative to perivascular, peripheral and infiltrating macrophages. Often called “homeostatic” genes. Example: *P2RY12*.
- Neutrophil, Monocyte: Elevated in neutrophils and monocytes relative to other types of myeloid cells. Example: *MMP8*.
- BrainMyeloid: This gene set contains the orthologues of the union of gene modules 2, 3, 5, 7 and 9 from ^8^ (column T of that manuscript’s Supplementary Dataset S4). These are genes elevated in resident relative to infiltrating and peripheral macrophages but not relative to perivascular macrophages. Example: *BIN1*.
- LPS-Specific: Genes that are significantly induced in myeloid cells by LPS but not significantly changed in myeloid cells in response to LCMV, β-amyloid, Tau pathology, or SOD1G93A. Example: *TNF*.
- Myeloid-AD-Up (respectively, Myeloid-AD-Down) are the genes up (respectively, down) in this study in human AD myeloid cells at *P* ≤ 0.05 and Max Cook’s P ≥ 0.01 (no fold-change cutoffs were applied). Example: *IL15* (respectively, *CECR2*).
- Barres-Neuron: Enriched in neurons relative to other CNS cell types from Ben Barres’ lab’s RNA-Seq profiles from fresh sorted cells from surgical resections ^4^. The specific genes are the same as those used in Fig. 1b and can be found in column N of Supplementary Dataset S5 from ^8^. Example: *SNAP25*.
- HumanMyeloid: This is a refinement of the “Myeloid” genes derived from the Barres data set ^4^. We felt that our starting list, in column N of Supplementary Dataset S5 from ^8^, included some genes that might have reflected activated microglia or other cell types, or that were not robustly expressed in multiple datasets. Therefore, from that list we excluded genes that were DE between perivascular macrophages and parenchymal microglia from an unpublished RNA-Seq study, or that were DE in brain myeloid compartment form LPS-treated (GSE75246) or PS2APP (GSE89482) animals. We also discarded genes that did not have median nRPKM ≥ 1 in the control myeloid cells from this study, or from a control group of an unpublished mouse RNA-Seq study. Finally, we removed NABP1 and OTUD1 because we thought based on visual inspection of their expression values that they would not be ideal markers. The final list is in column P of this manuscript’s Supplementary Table 2. Example: *ITGAM*.

Gene set scores (Fig. 2c, 2d, 3a and Extended Data Figs. 3a, 4) were calculated as described ^8^. Briefly, gene expression values were first log-transformed and stabilized as Log2(nRPKM+1), or, for ROSMAP-DLPFC, Log2(normalized RSEM+1). Then the average log-scale expression values of the controls were subtracted out for each dataset to yield control-centered gene expression values. The gene set score for a sample was then calculated as the average over all genes in the set of the control-centered gene expression values. For DE scores (Fig. 2c and Extended Data Figs 3a, 4a) a similar method was used, but with a signed average: up genes were weighted by +1 and down genes by −1.

Myeloid balancing was performed as described ^8^, but using the HumanMyeloid (described above) rather than complete Barres-Myeloid set of myeloid markers.

We also cross-checked specific mouse studies for potential insights, but we saw little overlap between our human DE genes and DE genes associated with PS2APP (GSE89482) or 5XFAD (GSE65067) beta-amyloid model microglia; Tau-P301S FTD model microglia (GSE93180); microglia following LPS or LCMV injection (GSE67858, GSE75246); old versus young mouse microglia (GSE62420); cerebellar versus cortical microglia (GSE62420); perivascular macrophages relative to parenchymal microglia (GSE60361); neutrophil versus other CD11b+ brain-resident cells (unpublished data); or infiltrating macrophages versus brain-resident microglia (GSE68376) ^6,8,10–12,29,30^ (analyses not shown).

**Supplementary Information** is linked to the online version of the paper at www.nature.com/nature.

## Acknowledgements

We wish to acknowledge Jeremy Stinson, Joe Guillory, Karen Toy and Subhra Chaudhuri for assistance with library preparation and sequencing; Chris Bohlen, Kevin Huang, Julia Kuhn, and Amy Easton for helpful critique of the manuscript; and Alison Goate for the suggestion to use the term “damage-associated microglia” instead of “disease-associated microglia”. We are grateful to the Banner Sun Health Research Institute Brain and Body Donation Program of Sun City, Arizona, for the provision of human brain tissue. The Brain and Body Donation Program has been supported by the National Institute of Neurological Disorders and Stroke (U24 NS072026 National Brain and Tissue Resource for Parkinson’s Disease and Related Disorders), the National Institute on Aging (P30 AG19610 Arizona Alzheimer’s Disease Core Center), the Arizona Department of Health Services (contract 211002, Arizona Alzheimer’s Research Center), the Arizona Biomedical Research Commission (contracts 4001, 0011, 05-901 and 1001 to the Arizona Parkinson’s Disease Consortium), and the Michael J. Fox Foundation for Parkinson’s Research.

## Author Contributions

KS, BAF and DVH conceived of the study and planned the analysis. KS performed experimental procedures for the sorted cell RNA-Seq (GSE125050; tissue handling, cell sorting, RNA extraction) and qPCR (Fig. 2b, c and Extended Data Fig. 3). KS and ZM’s group performed RNA-Seq library construction, and ZM oversaw library QC and sequencing. AE, MPvdB and OD generated whole tissue RNA-Seq data, and MAH performed initial data processing of the new FuG bulk tissue study (GSE125583). BAF performed all other data analysis and generated all figures except for Extended Data Fig. 1. TGB and GES performed human subject and tissue selection and reviewed the manuscript. BAF and DVH wrote the manuscript.

## Author Information

Reprints and permissions information is available at www.nature.com/reprints. TGB and GES participated in this study under a contracted research agreement. All other authors are current or former employees of the pharmaceutical company Genentech, Inc., and declare no further competing interests. Correspondence and requests for materials should be addressed to.

